# Dynamism and context dependency in the diversification of the megadiverse plant genus *Solanum* L. (Solanaceae)

**DOI:** 10.1101/348961

**Authors:** Susy Echeverrĩa-Londoño, Tiina Särkinen, Isabel S Fenton, Sandra Knapp, Andy Purvis

**Author notes:** Corresponding author: Susy Echeverrĩa-Londoñ; Email.

## Abstract

- Explosive radiations have been considered one of the most intriguing diversification patterns across the Tree of Life, but the subsequent change, movement and extinction of the constituent species makes radiations hard to discern or understand as geological time passes.
- We synthesised phylogenetic and distributional data for an ongoing radiation — the mega-diverse plant genus *Solanum* L. — to show how dispersal events and past climatic changes have interacted to shape diversification.
- We found that despite the vast diversity of *Solanum* lineages in the Neotropics, lineages in the Old World are diversifying more rapidly. This recent explosive diversification coincides with a long-distance dispersal event from the Neotropics, at the time when, and to places where, major climatic changes took place. Two different groups of *Solanum* have migrated and established in Australia, but only the arid-adapted lineages experienced significant increases in their diversification, which is consistent with adaptation to the continent’s long-term climatic trend and the diversification of other arid-adapted groups.
- Our findings provide a clear example of how successful colonisation of new areas and niches can – but do not always – drive explosive radiations.

## Introduction

The uneven distribution of taxonomic diversity among the branches of the tree of life and geographic regions is one of the most intriguing puzzles in biology. The increasing availability of global field study data and large phylogenies continues to reveal how some groups are highly diverse compared with their depauperate sister groups. Even though we expect the stochastic nature of the diversification process to produce asymmetric patterns of diversity which do not necessarily require deterministic explanations (Raup *et al.*, 1973; Slowinski & Guyer, 1993; Purvis, 1996), identifying significant patterns of unequal diversity (i.e., those that depart from the null expectation) throughout the tree of life has the potential to reveal important mechanism about the origin and maintenance of biodiversity.

One of the main hypotheses in macroevolution is that unequal diversity can arise from significant differences in net diversification rates (i.e., speciation – extinction) among lineages and through time and space (Stanley *et al.*, 1981; Alfaro *et al.*, 2009; Wiens, 2011, 2015). This asymmetry in evolutionary rates is often interpreted as resulting from the presence of a spectrum of ecological opportunities, such as the opening of niche space by the development of new key traits, extinction of antagonists, and/or colonisation of new open areas (Simpson, 1955; Schluter, 2000; Moore & Donoghue, 2007; Losos, 2010; Yoder *et al.*, 2010; Rabosky *et al.*, 2013). The first step to uncover the relative influence of these factors in diversification is to decompose the macroevolutionary dynamics of a clade into its essential components ‐‐‐speciation and extinction rates. Phylogenetic Comparative Methods (PCM) provide a series of approaches to quantifying these rates using reconstructed phylogenies along with various stochastic models (see Stadler, 2013 and Morlon, 2014 for a detailed review of diversification approaches). However, the results of such approaches are often limited and challenged by the difficulty of accounting for extinct species usually not included in molecular phylogenies; these factors are especially problematic in older clades (Liow *et al.*, 2010; Quental & Marshall, 2010; Rabosky, 2010). Furthermore, poor species-level sampling and branch support in the underlying phylogenies used for testing these hypotheses usually limits the power of tests of diversification dynamics. Therefore, a focus on recently emerged clades that are species-rich, widespread and are underpinned by well-sampled and robust phylogenies will provide us with a more accurate picture about the macroevolutionary dynamics shaping biodiversity.

In this study, we investigate the tempo and mode of species diversification of the mega-diverse plant genus *Solanum* L. (Solanaceae). This genus is one of the most diverse and economically important plant groups with around 1,500 species distributed worldwide (i.e., almost the half of the diversity of the Solanaceae family). *Solanum* contains species from all major tropical and subtropical biomes and shows high heterogeneity of richness among its subclades. This plant genus has been traditionally divided into two major groups, the spiny (prickly species) and non-spiny solanums. With robust monophyletic support, the spiny solanums (known as the Leptostemonum clade, or subgenus *Leptostemonum Bitter*) include ca. 420 species and can be recognised by the presence of stellate trichomes, prickles, and long tapering anthers (Vorontsova *et al.*, 2013). Spiny solanums are most diverse in the Neotropics (ca. 150 species), but also have high diversity in Australia (ca. 130) and Africa (ca. 79). This high diversity of spiny solanums out of the Americas contrasts with the distribution of the diversity of the nonmonophyletic group — the non-spiny solanums — which has more than 90% of its ca. 670 species distributed in the Neotropics.

As one of the top ten species-rich genera of angiosperms (Frodin, 2004)*, Solanum* is suggested to have high diversification rates, especially in the Neotropics which contains most of its diversity and endemism (Olmstead & Palmer, 1997; Särkinen *et al.*, 2013; Knapp & Vorontsova, 2016). However, where radiating lineages occur and how much they contribute to current diversity and geographic differences in richness among subclades of *Solanum* has not yet been explored. Here, we assembled the first set of complete time-calibrated and species-level phylogenies of extant *Solanum* species (1169 species), to reconstruct diversification rates across lineages and analyse them in a geographical context. We test the origins of the high heterogeneity of species richness observed among subclades of *Solanum*, investigating the relative importance of clade-specific, tree-wide and geographic variation in evolutionary rates, as well as how these patterns are associated with historical biogeographic movements and/or environmental changes.

## Materials and Methods

Phylogenetic relationships and divergence times among the main clades of *Solanum* were obtained from the Särkinen *et al.*, 2013 Solanaceae phylogeny. This tree was built using two nuclear and six plastid loci from 454 *Solanum* species (ca. 34% of the extant species). This phylogeny identified a total of 37 clades and subclades within the genus, with a vast heterogeneity of species richness within these.

### Correcting for non-random taxon sampling

The incomplete nature of the Särkinen *et al.*, 2013 phylogeny could be problematic because non-random incomplete sampling in phylogenetic studies could significantly affect the estimate of macroevolutionary rates (Höhna *et al.*, 2011). Despite the claims that current speciation-extinction models account for incomplete species sampling (e.g., MEDUSA, Alfaro *et al.*, 2009; BAMM, Rabosky, 2014; TreePar, Stadler, 2011; TESS, Höhna *et al.*, 2016b; RevBayes, Höhna *et al.*, 2016a), these models often assume random, even sampling of the species across the phylogeny (i.e., each terminal in the phylogeny is assumed to have the same probability of being sampled). However, this assumption is very often violated in empirical examples due to geographic, temporal and taxonomic sampling biases. To account for non-random incomplete taxon sampling in the diversification analysis, we used the stochastic polytomy resolver PASTIS (Thomas *et al.*, 2013), to place missing species of *Solanum* in the phylogeny using taxonomic constraints following a birth-death model. This method is a conservative approach for increasing the sampling in phylogenies since it infers the timing of missing splits under a constant rates birth-death model. Therefore, any inference of rate-heterogeneity in the final analysis will have stronger support since this approach is expected to bias towards the detection of constant-rate models of diversification (Thomas *et al.*, 2013). Every extant and accepted *Solanum* species was assigned to one of the clades (listed in Supporting Information Table S1) using the Särkinen *et al.*, 2013 phylogeny as a backbone. Complete trees were then generated for each clade of *Solanum* using a combination of species with genetic data and taxonomic constraints. For each clade, PASTIS creates an output file in a nexus format, which contains the full set of tree constraints ready to be executed in MrBayes. Posterior distributions of phylogenies for each clade were then inferred in MrBayes 3.2.3 (Ronquist & Huelsenbeck, 2003) using a relaxed clock model (independent branch rates – igr prior), with the default (exponential) prior on the distribution of branching rates. A distribution of 100 dated trees was produced containing nearly all described species of *Solanum*, 1169 species in total (excluding those with uncertain taxonomic position), after sampling and grafting the clades distributions. For further information see Supporting Information Methods S1.

### Diversification analysis

Several approaches have been developed to detect significant shifts in diversification across the branches of a phylogeny. In this analysis, we mainly used the BAMM approach (Rabosky, 2014), a Bayesian framework for reconstructing evolutionary dynamics from phylogenetic trees, which aims to provide an improvement on methods that identify heterogeneity in rates only across specific subclades, such as MEDUSA (Alfaro *et al.*, 2009). However, other approaches such as TESS (Höhna *et al.*, 2016b) and RevBayes (Höhna *et al.*, 2016a), were also implemented to assess the robustness of the results obtained in BAMM.

The underlying branching process in BAMM includes the effect of time-dependence on diversification (i.e., the age of a lineage can affect its diversification rate). It also implements a diversity-dependent diversification model where the number of lineages in a clade may affect its diversification rates. BAMM uses a reversible-jump Markov Chain Monte Carlo approach to explore a larger space of parameters and candidate models of diversification. Since it follows a Bayesian statistical framework (rather than the maximum likelihood framework used in MEDUSA), BAMM implicitly accounts for the uncertainty in parameter estimates by providing a distribution of marginal posterior probabilities instead of point estimates. To implement this approach in the *Solanum* phylogeny, we used the distribution of 100 complete species trees (generated using PASTIS, see previous section), to consider the phylogenetic uncertainty in the estimates of diversification rates. We set the priors of speciation, extinction and the expected number of diversification shifts (g=1) using the R package BAMMtools v 2.0.5 (Rabosky *et al.*, 2014), which identifies the priors of the diversification parameters based on the distribution of divergence times of the phylogeny. For each of the 100 trees, an MCMC analysis was performed with four separate runs of 20 million generations. All the analyses were run using the C ++ BAMM command line program v 2.5.0 on the Imperial College London’s High-Performance computing cluster (http://www.imperial.ac.uk/admin-services/ict/self-service/research-support/hpc/). We then checked for convergence of the MCMC samples making sure the effect sample size was at least 200 for both the number of evolutionary shifts and likelihood using the CODA R package V 0.16-1 (Plummer *et al.*, 2006). The first 25% of the samples were discarded as burn-in.

For each of the 100 trees run in BAMM, a distribution of 1000 samples of the posterior probabilities of diversification were created. Each sample from the posterior includes either a single event (i.e., the diversification is described by a single time-varying process — no shifts in diversification) or a mixture of one or more shifts and associated parameters. For each set of posterior probabilities, we extracted the list of nodes associated with “core” rate shifts (i.e., rate shifts with a marginal probability significantly higher than the probability expected from the prior alone) and calculated the frequency with which these nodes are associated with significant rate shifts across the 100 trees run in BAMM (i.e., across a distribution of 100 × 1000 trees). Finally, we computed the mean of diversification rates for each of the species present in the Särkinen *et al.*, 2013 phylogeny across the pooled distribution of posterior samples produced in the BAMM analysis.

Studies such as Moore *et al.*, 2016 have raised concerns about several methodological issues in BAMM including a potentially incorrect likelihood function, the strong influence of priors on posterior estimates, and some theoretical errors (e.g., the incorrect use of the Poisson distribution as the error distribution of the prior number of shifts of diversification). To assess the effects of these potential limitations on the results, we performed several sensitivity analyses focusing on the two most important issues stated by Moore et al., 2016, (1) the prior sensitivity of posterior distribution on the number of rate shifts, and (2) the reliability of the diversification rate estimates. We performed these analyses using alternative approaches such as TESS (Höhna *et al.*, 2016b) and RevBayes (Höhna *et al.*, 2016a). Further details about the methodology and results of these analyses can be found in Supporting Information Methods S2.

### Geographical patterns of diversification

Using 64,826 unique records from 1,096 taxa (see Supporting Information Methods S2 for further details), we mapped the mean assemblage diversification rates of *Solanum* on a 1 × 1 degree map to determine which regions have accumulated or are currently accumulating a higher or lower number of *Solanum* lineages across the globe. The mean assemblage diversification rate was calculated as the geometric mean of all species-specific rates present in a grid cell. We also computed a weighted version of this, dividing the mean species-specific diversification rates by the inverse of their range size ‐‐‐ log of the area (m^2^) occupied by each species, to correct for the diversification rates in areas with a high occurrence of widespread species (i.e., large ranging species have less weight in the overall diversification rates of a grid cell).

Finally, we reconstructed lineages-through-time plots for the three-principal diversity centres for *Solanum* (Neotropics, Australia and Africa) using the R package “paleotree” v 2.3 (Bapst, 2012) to visualise the geographical differences in lineage accumulation dynamics among continents.

### Ancestral range reconstruction

We investigated the historical biogeography of *Solanum* using the R package “BioGeoBEARS” (Matzke, 2012, 2016) — BioGeography with Bayesian (and likelihood) Evolutionary Analysis in R Scripts. This approach provides a statistical framework to compare traditional models in biogeography such as DIVA (Dispersal-Vicariance Analysis, Ronquist, 1997), DEC (Dispersal-Extinction-Cladogenesis, Ree *et al.*, 2005), and BayArea (Landis *et al.*, 2013). Each of these models assume different processes to reconstruct the ancestral range of lineages. In all the models, species ranges are allowed to change along the branches by anagenetic evolution through two main events: dispersion (range expansion) and extinction (range contraction). The events allowed in cladogenesis vary depending on the fitted model. For example, sympatric speciation subset, or peripatric speciation (i.e., the ancestral range is completely inherited by one of the daughters whereas the other inherits only a subset of the range) is only allowed in the DEC model. However, unlike the DIVA and the BayArea models, the DEC model only allows one daughter to inherent widespread distributions during a vicariant event, since the widespread distribution of both daughters would assume additional events such as post-speciation dispersal (Ree *et al.*, 2005). In addition to these classic events of biogeography, BioGeoBEARS includes founder-event speciation (+j) which considers the influence of speciation through long-distance dispersal, such as is common in Island-like models.

To avoid the effects of birth–death polytomy resolvers on the natural patterns of trait phylogenetic structure (following Rabosky, 2015), we reconstructed the ancestral ranges of *Solanum* using the time-calibrated clade credibility (MCC) from Särkinen *et al.*, 2013. Several species were pruned from the tree as follows: (1) species which are considered widely cultivated or with ambiguous native distribution, (2) species considered as synonyms, and therefore duplicated in the tree, and (3) species with a low support of a direct ancestor-descendant relationships leading to negative branches in the MCC. The final pruned phylogeny used for subsequent analyses contained 386 species of *Solanum* with the addition of *Jaltomata andersonii* as the outgroup.

Using the extracted distribution records, we created an occurrence matrix of *Solanum* species split into six biogeographic areas—Africa, Australia, Indo-Pacific, Neotropics, Nearctic and Palearctic based on the floral kingdoms defined by Cox (2001). The distribution of widespread species was double checked and corrected for potentially recent cultivated and/or naturalised species using the native descriptions available from http://solanaceaesource.org or specific taxonomic monographs. The maximum number of areas in the ancestral range reconstruction was set to three to reduce the complexity and computational time in the analysis.

Three biogeographic models were fitted to the *Solanum* phylogeny and the associated geographic distributions using BioGeoBEARS — DIVALIKE, DEC and BAYAREALIKE models. The influence of founder-event speciation event (+j) was also included into each model, resulting in a total of six models. An additional set of six biogeographic models were fitted using a dispersal matrix multiplier to weight the dispersal probability of adjacent areas as 1, 0.5 and 0.001 for easy, medium, and hard dispersal. The model that best describes the empirical data (i.e., optimal fixed model structure) was then chosen using a stepwise selection from the candidate models ranking under the Akaike Information Criterion (AIC, Burnham & Anderson, 2002).

Once the best-fitting model was identified, we estimated the overall probabilities of the anagenetic and cladogenetic events conditional on the model, the phylogeny, and the geographic distributions from 100 Biogeographic Stochastic Maps (BSM) to account for the uncertainty in the state transitions and ancestral range reconstructions (Matzke, 2016). These stochastic maps are similar to simulations of trait change along phylogenies (Huelsenbeck *et al.*, 2003) using transition rate models (Pagel, 1999). Given the observed range data, the phylogeny and the best fitting model of biogeographic events, the BSM simulates possible histories constrained by the observed ancestral ranges. The ancestral state probabilities obtained under the best fitting model are equivalent to the average of all the probabilities of the simulated histories from the biogeographic stochastic maps (Matzke, 2016).

## Results

We found significant heterogeneity in diversification rates along the branches of the *Solanum* phylogeny. This variation in evolutionary rates has contributed to the pronounced disparities in species richness among groups; the Old World spiny clade and the Petota subclade are supported as the most rapid radiations (Fig. 1). The Old World clade has diversified nearly as twice as rapidly as any other group, with approximately 0.68 lineages per million years (lineages Myr^−1^) as compared to the global mean of 0.25 lineages Myr^−1^ (Fig. 1). The node supporting the crown group of the Old World clade was the node with the highest frequency of shifts found across the 100 runs. Other groups such as the Petota subclade (from the Potato clade, Petota + Tomato) and the node supporting the Leptostemonum group (i.e., the spiny solanums) also show some signal of shifts in diversification but with a weaker support compared with the Old World clade (Fig. 1, see also Supporting Information Fig. S2, S3).

**Fig. 1.**
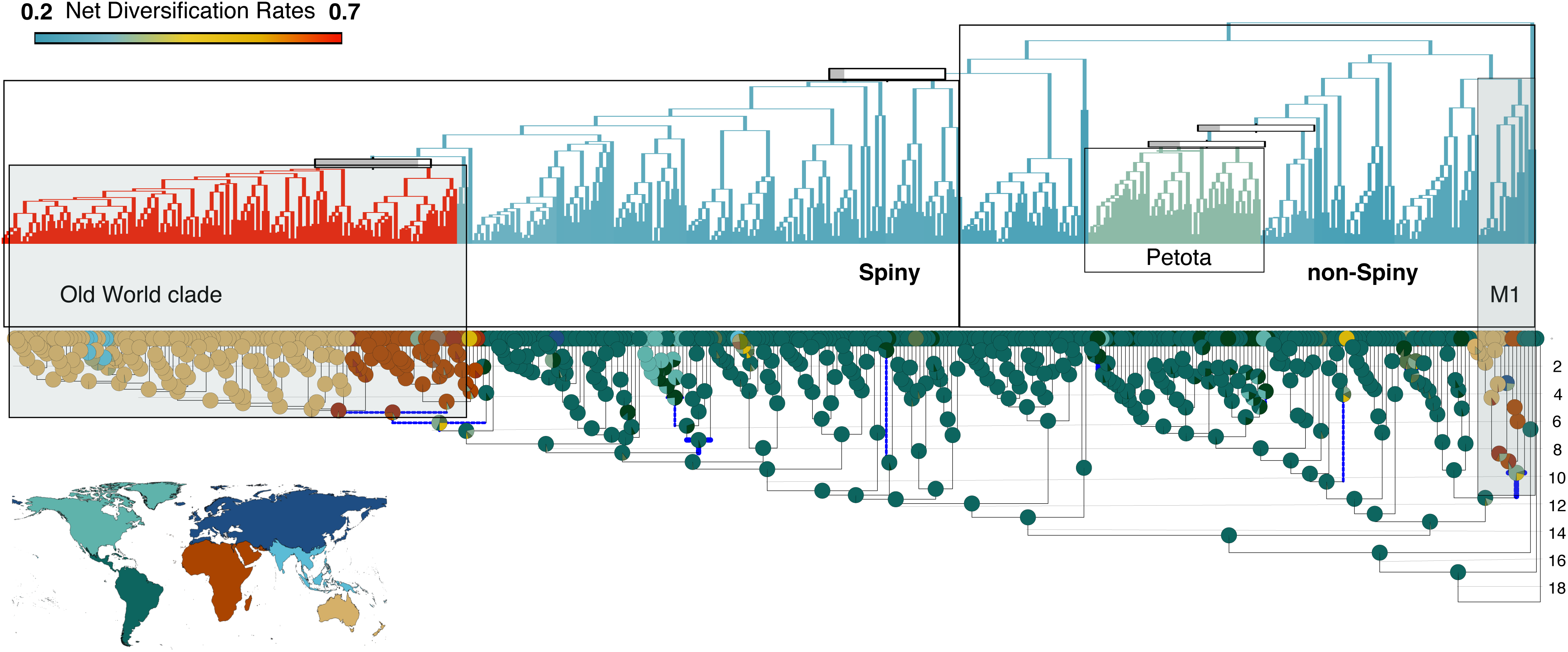
Mean net diversification rates and historical biogeographic reconstruction across the phylogeny of *Solanum*. For the net diversification rates, bars represent the frequency with which nodes are associated with shifts in diversification rates across a distribution of 100 complete phylogenies of *Solanum* (see Methods for further detail). For the historical biogeographic reconstruction, pies at each node represent the probabilities of the ancestral range distribution; the map indicates the regional colour code. The branches highlighted in blue represent the dispersals events inferred in at least the 50% of Biogeographic Stochastic Maps (BSMs). The timescale is given in Myr.

The number of diversification shifts found in the BAMM analysis is largely robust to the defined prior distribution. Supporting Information Fig. S7-S10 show that across the 100 *Solanum* phylogenies run in BAMM, the distribution of estimated shifts (posterior samples) differs significantly from the distribution of the expected number of shifts (prior). Although the prior distribution (γ=1) applied in this analysis was strong and conservative (i.e., the zero-shift model was set to be the most likely outcome), the zero-shift model was never sampled in the posterior for any of the BAMM runs showing overwhelming evidence of the heterogeneity of shifts found in this analysis. Moreover, the sensitivity analysis based on a subset of the data (i.e. the Särkinen et al. 2013 phylogeny without the additional species) found that the number of shifts was not sensitive to different priors of diversification rates (i.e., different values of γ, 0.5,1,2,10,100) as shown in Supporting Information Fig. S11. Additionally, the analysis of diversification rates in RevBayes showed similar results to those from the BAMM analysis (see Supporting Information Fig. S3, S4).

Although most of the diversity and endemism of *Solanum* is found in the Neotropics (ca. 70% of species, see Fig.1, 2a), the highest diversification rates are seen in lineages mainly concentrated in Australia, Africa and the Indo-Pacific (Fig. 2b). Africa shows a heterogeneity of rates as it contains species from groups with standard diversification rates such as Morelloids (0.27 lineages Myr^−1^) and African non-spiny (0.25 lineages Myr^−1^) as well as species from the rapidly diversifying Old World clade (0.68 lineages Myr^−1^).

**Fig. 2.**
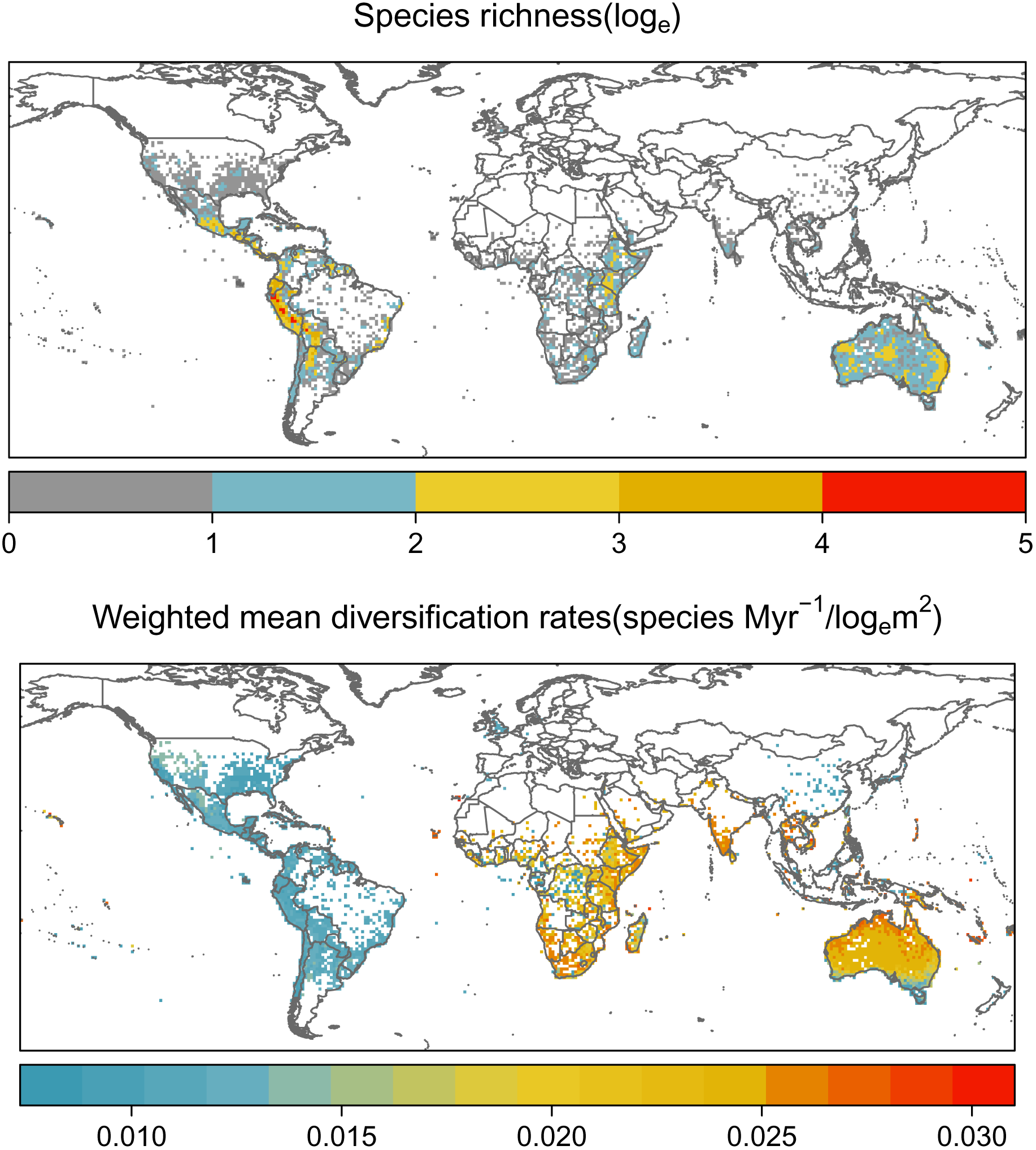
Global distribution of species richness (a) and the net diversification rates of *Solanum* (b) using 1 × 1 degree grid cells. Weighted mean lineage net diversification rates were calculated as the mean net diversification rate for all the species present in a grid cell assemblage, weighted by the inverse of their range size, *log_e_m^2^*

Lineages-through-time curves (Fig. 3a) confirm that most of the diversification in *Solanum* has occurred within the Neotropics. At the global level, there is a slight acceleration in the number of lineages of *Solanum* in the last 5 Myr, which is likely to be shaped by the considerable increase of lineages in Australia around the same time. Australia shows an interesting latitudinal heterogeneity of diversification rates (Fig. 3b) shaped by the distribution of lineages of different evolutionary origins — species from the Archaesolanum subclade, which belongs to the M1 clade in Fig. 1, are mainly distributed in the south with 0.20 lineages Myr^−1^ whereas the Old World spiny species with 0.68 lineages Myr^−1^ have a widespread distribution.

**Fig. 3.**
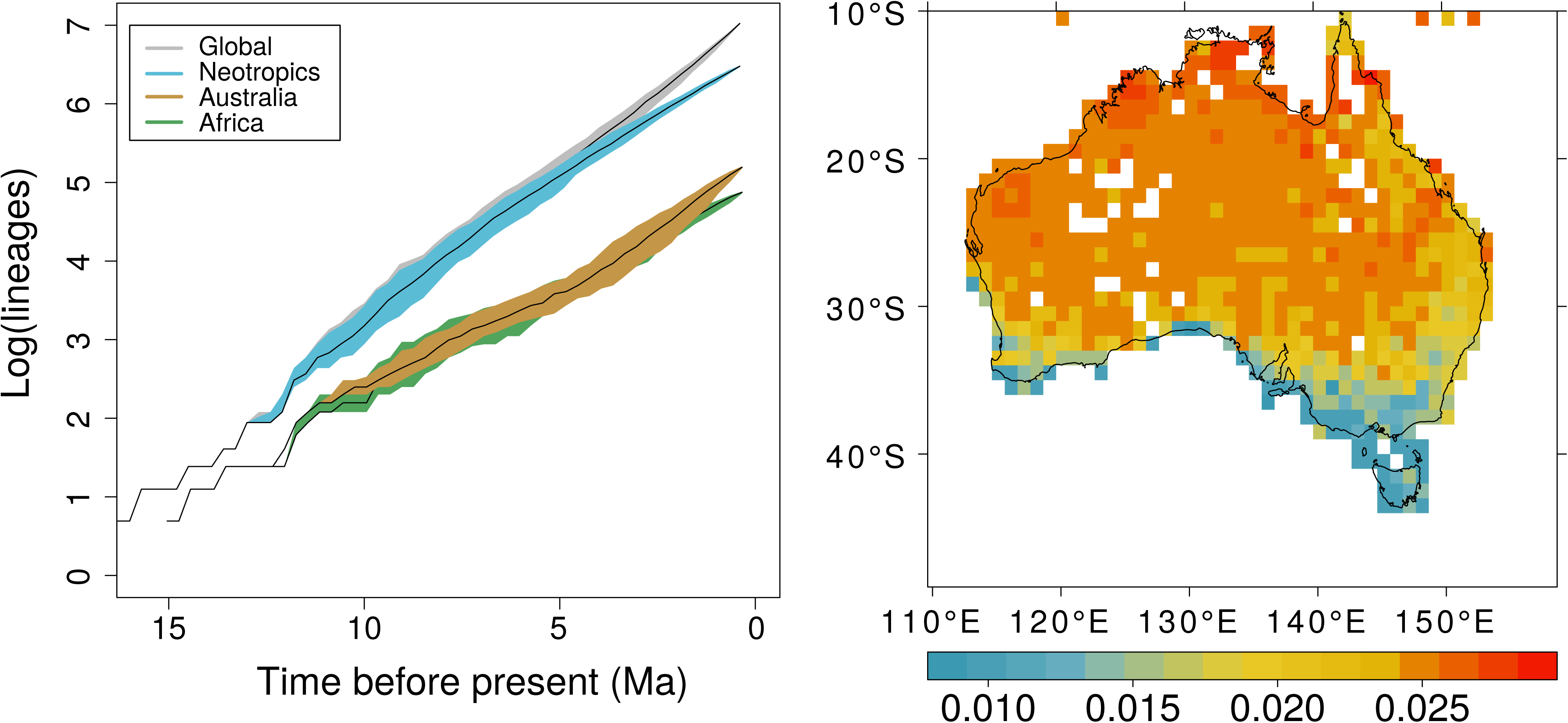
(a) Lineages-through-time plot of *Solanum* species for the global and three regional diversities (Neotropics, Australia, and Africa). Coloured areas around the median curves, shown in black, represent two-tailed 95% upper and lower quantiles of the number of lineages across a distribution of 100 complete phylogenies of *Solanum* (see Methods for further details). (b) Distribution of the net diversification rates of *Solanum* in Australia (same units as in Fig. 2b).

Ancestral range reconstructions across the *Solanum* phylogeny are best explained by the DEC M1 model (Dispersion, Extinction and cladogenesis), which allows for equal probabilities of dispersion from the Neotropics to both Africa and Australia (Supporting Information Table S2). Including founder-event speciation in the model did not improve the AIC values significantly (∆ AIC = 0.5), therefore, all the historical biogeographic results, including the Biogeographic Stochastic Mapping, in this study were based on the simpler model, DEC M1.

The genus *Solanum* appears to have risen in the Neotropics ca. 15 Myr (95% HPD 13-18), as do the majority of its subclades (see Fig. 1). The distribution of *Solanum* appears to have occurred mainly via within-area sympatric speciation and dispersion, with vicariance only supported in 3% of the total events (see Supporting Information Table S3). The distribution of *Solanum* in the Nearctic region occurred through at least seven dispersal events out of the Neotropics. Although several subclades are currently distributed in the Old World, only two subclades appear to have arisen there via historical events (as opposed to recent introductions). These dispersal events from the Neotropics to the Old World occurred at different times and in two different groups of *Solanum*. The first movement from the Neotropics to Africa + Australia is likely to have occurred ca. 10 Myr (95% HPD 7-12) within the non-spiny solanums, in the crown group of the subclades *Solanum valdiviense* + African-non-spiny + Normania + Archaesolanum (referred to as the M1 subclade on Fig. 1). The second dispersal event occurred ca. 6 Myr (95% HPD 5-7) within the spiny solanums in the subclade Elaeagnifolium + Old World spiny clade (see Fig. 1). The direction of these dispersal events from the Neotropics to either Africa and Australia is still unresolved from the stochastic mapping estimates in both the non-spiny solanums (a 30% dispersal probability from the Neotropics to Africa, and 24% to Australia) and the spiny solanums (a probability of 45% of dispersing from the Neotropics to Africa, and 29% to Australia).

Overall, the Neotropics were the main source of *Solanum* movements with more than 60% of the estimated dispersal events (Supporting Information Fig. S12, S13). The most frequent dispersals involved movements from the Neotropics to the Nearctic with ca. 40% of the total estimated events. Movements from the Neotropics to Australia appear to be more frequent than those from the Neotropics to Africa (8.7 ± 1.1 vs 2.9 ± 0.9). Dispersals from Africa to Australia are more common than those in the opposite direction (5.5 ± 1.2 vs 1.64 ± 0.9).

## Discussion

In this study, we demonstrate that the great diversity of *Solanum* is likely to be the result of a great heterogeneity in evolutionary rates among its subclades supported by at least two recent radiations — the Old World spiny clade and the Petota subclade (Fig. 1). Previous studies such as Whalen, 1979 and Whalen & Caruso, 1983 were unable to identify unusual rates of speciation within *Solanum*, concluding that the great taxonomic diversity of this genus may reflect a gradual accumulation of species in a relatively old clade (which at the time was believed to have a Cretaceous origin, Hawkes & Smith, 1965). However, the results of these previous studies were based on species-poor but well-known sections of *Solanum* such as Lasiocarpa (13 species) and Androceras (12 species) which ignored the great heterogeneity of evolutionary regimes present in other groups of the genus. Our results demonstrate the importance of accounting for among-clade rate heterogeneity in understanding the complex dynamics of diversification of megadiverse groups such as *Solanum* (Ricklefs, 2007; Morlon, 2014).

Despite most of the richness and endemism of *Solanum* occurring in the Neotropics (Fig. 2a), this region has lower diversification rates than the Old World (Fig. 2b). This mismatch between the high diversity of *Solanum* within the Neotropics and its net diversification rates could be explained by the early and long history of diversification of *Solanum* in the Neotropics. *Solanum* lineages within the Neotropics are older and, therefore, had more time and opportunity to accumulate before establishing in other regions, creating the current imbalance of diversity (McPeek & Brown, 2007; Weir & Schluter, 2007). The Neotropics is the region where *Solanum* species experienced the greater within-area speciation events, which has also been demonstrated as the highest within the family Solanaceae as a whole (Dupin *et al.*, 2016). The long history of diversification and expansion of *Solanum* species within the Neotropics, therefore, drove a greater lineage accumulation compared with other regions (Fig. 3a), making the Neotropics the main source of diversity and origin of biogeographic movements of solanums across the globe (Fig. 1, and Supporting Information Fig. S12, S13).

There are no obvious novel morphological or physiological traits associated with the diversification of the Old World clade, given this group is not well defined by any single distinctive morphological traits or combination of morphological characters (Stern *et al.*, 2011), but instead is defined by its geographical distribution and robust monophyly (Levin *et al.*, 2006; Weese & Bohs, 2007, 2010; Stern *et al.*, 2011; although Aubriot *et al.*, 2016 showed that six species thought to belong to the Old World clade, based on their distribution, now belong to the Torva clade). However, the signal of change in diversification at the base of the clade supporting all spiny solanums in Fig. 1, indicate that the distinctive traits of spiny solanums as a whole, such as stellate indumentum and prickles (Whalen, 1984) could have played an essential role in the diversification of the Old World clade. In this context, the Old World clade could be an example of an exaptive radiation where previously acquired traits – originally shaped by different selective forces – are advantageous under a new selective regime (Simões *et al.*, 2016). Furthermore, this nested radiation of lineages of the Old World clade within spiny solanums might be a classic example of Key confluence *sensu* Donoghue & Sanderson, 2015, where shifts in diversification are associated with the interaction between key innovations and extrinsic factors such as biogeographic movements and/or environmental changes, as shown in our ancestral range reconstruction (Fig. 1).

We found that not all the *Solanum* groups distributed in the Old World (i.e., mainly in Africa, Australia, and the Indo-Pacific) that were associated with long-dispersal events resulted in explosive radiations such as occurred in the Old World clade. This suggests that distinct ecological opportunities and evolutionary regimes might have shaped the diversity of spiny and non-spiny *Solanum* in the Old World. The lineages of non-spiny solanums present in the Old World are much less diverse than the spiny solanums and occur in different environments. In Africa, the non-spiny solanums are less diverse despite being significantly older than the spiny solanums; they are mostly found in forested and mesic regions (Knapp & Vorontsova, 2016). In Australia, the diversity of non-spiny solanums is represented by the Archaesolanum group known as the Kangaroo apples. With eight species in total (Poczai *et al.*, 2011), this group occurs in the temperate and forested areas of the South West Pacific (e.g., Australia, Tasmania, New Zealand and Papua New Guinea). In contrast, spiny solanums occurring in continental Africa and Australia are much more diverse and are predominantly concentrated in arid areas (Symon & Others, 1981; Whalen, 1984; Bean, 2004; Vorontsova *et al.*, 2013). In Africa, these are in vegetation types such as the “Somalia-Masai regional centre of endemism” *sensu* White (1983), whilst in Australia, the ca. 200 species of spiny solanums are predominantly distributed in warmer and arid regions (Fig. 3b).

A comparison of temporal patterns of lineage accumulation among major regions shows a distinctive signature of a rapid diversification within Australia at ca. 5 Myr (Fig. 3a). The timing of diversification and dispersion of spiny solanums in the Old World, as well as their concentration in arid and semi-arid regions of Australia (Fig. 3b), suggests a potential link between past environmental changes and radiation of this group. A similar signature has been shown in a wide range of groups from Australia (Harmon *et al.*, 2003; Crisp *et al.*, 2004; Crayn *et al.*, 2006; Crisp & Cook, 2007, 2013; Rabosky *et al.*, 2007; Byrne *et al.*, 2008, 2011; Harrington *et al.*, 2012; Blom *et al.*, 2016). The dramatic climate change in Australia since its separation from Gondwana at ca. 50 Myr is likely to have triggered the radiation of several taxa (Byrne *et al.*, 2008) and the extinction of others (Byrne *et al.*, 2011). The initial stage of aridification in the mid-Miocene (ca. 15 Myr) and the late expansion of desert regions of Australia in the last 1-4 Myr, has led to the assembly of new biomes, such as the arid, monsoonal and alpine regions, encouraging the establishment and expansion of arid-adapted lineages (e.g., Crisp *et al.*, 2004; Crayn *et al.*, 2006; Rabosky *et al.*, 2007; Byrne *et al.*, 2008; Crisp & Cook, 2013). This pattern and the congruence of the timing of diversification with other arid-adapted Australian groups could indicate that the Old World spiny solanums established and diversified in Australia as the arid environments expanded. The widespread distribution of Old World spiny solanums in Australia suggests a positive effect of distribution area on diversification rates. The large expanse of arid habitat in Australia may have created opportunities for allopatric speciation within the biome itself (Losos & Schluter, 2000; Davies *et al.*, 2005; Rabosky *et al.*, 2007). The successful expansion and diversification of spiny solanums in warm and arid zones in Australia, in contrast with the slower diversification of the more mesic non-spiny clades, could indicate the influence of features adapted to warm and dry environments on species diversification. Moreover, the sister group of the diverse Old World clade is the species-poor Elaeagnifolium clade, with five species confined to semi-arid regions of North and South America (Knapp & Vorontsova, 2016), suggesting that some pre-adaptation traits to dry conditions may have driven diversification in the Old World. This evidence suggests that the dispersal of spiny solanums to the Old World, the influence of past environmental changes, as well as the preadaptive condition of spiny solanums to colonise warm and arid regions contributed to the explosive radiation of *Solanum* lineages in Africa and Australia.

## Implications of the study

The dynamics shaping the diversity of species-rich mega-diverse genera may be more complex than previously thought. A variety of mutually reinforcing drivers appear to have formed the current patterns of diversity in *Solanum*, which cautions against single-cause hypotheses, such as the influence of “key innovations” often invoked in macroevolutionary studies (Donoghue & Sanderson, 2015). These findings also raise important questions about the reliability of global studies that assume a single model for the evolution of character change and diversification across large phylogenies (e.g., Zanne *et al.*, 2014), or studies that use average values of diversity, trait distribution, or geographical distribution at generic level (e.g., Cornwell *et al.*, 2014). For instance, Beaulieu *et al.*, 2013 and Chira & Thomas, 2016 have demonstrated that heterogeneity of diversification rates can have significant consequences for the model selection of trait change that best fits comparative data. In general, therefore, the understanding of the magnitude and location of shifts in diversification rates should be included as prior information in macroevolutionary analyses, especially in analyses of trait evolution (Morlon, 2014; Chira & Thomas, 2016), or at least models of trait evolution must not implicitly assume that such shifts do not occur.

The results of this study also highlight the importance of context dependency in the study of diversification dynamics (De Queiroz, 2002; Donoghue, 2005; Donoghue & Sanderson, 2015). Our study shows that both opportunity and success in geographic movement, as well as correct environmental setting and climatic adaptations, were necessary to spark faster diversification in the spiny clade of *Solanum* in the Old World. Thus, focusing on effects of the combination of events and attributes on changes in diversification could represent a more productive framework in which to understand diversity processes (Donoghue & Sanderson, 2015). This combination could also explain why, although many correlates of diversification rates have been reported in the literature, few have demonstrated either generality or high explanatory power. Our results suggest the necessity of considering more complex analyses that involve the integration of phylogenies with other biological and Earth system data sources such as geography, climate, historical biogeography, and physiology.

We demonstrate that *Solanum* is a remarkable study system for understanding the diversification of plants, not only because it contains a recent and ongoing radiation, but also because its diversity is supported by several clades with different and sometimes contrasting evolutionary dynamics. There are still many drivers that need to be explored before we have the whole picture of the diversification of *Solanum*, especially correlations with biogeographic events and morphological traits, but increased taxonomic, phylogenetic, and morphological understanding of *Solanum* now means that we are beginning to uncover many aspects of the macroevolution of this mega-diverse plant genus, and can use it as a model for studying diversity in widespread, species-rich groups in general.

## Acknowledgements

This study was supported by the National Science Foundation (NSF) grant (to SK) ‘PBI *Solanum* – a world treatment’ DEB-0316614, and Colciencias (Administrative department of Science, Technology and Innovation of Colombia) who partially funded S.E.-L.’s PhD programme.

## Author Contribution

S.E.-L designed the study, analysed the data and wrote the first draft of the manuscript. TS provided the phylogenetic data (reconstructed the phylogeny), distribution and taxonomic data. ISF provided advice on the analysis and contributed to the main discussion of the macroevolutionary analyses. SK provided the taxonomic, distribution and diversity data, and supervised the study. AP designed and supervised the analysis. All authors reviewed the manuscript.

